# A cosmic view of ‘tundra gardens’: satellite imagery provides a landscape-scale perspective of Arctic fox ecosystem engineering

**DOI:** 10.1101/2022.08.23.504941

**Authors:** Sean M. Johnson-Bice, James D. Roth, John H. Markham

## Abstract

Animal ecology has benefitted greatly from advancements in remote sensing technology and data availability in recent decades. Most animal ecology studies using remote sensing data have focused on assessing how environmental characteristics shape animal abundance, distribution, or behavior. But the growing availability of high-resolution remote sensing data offers new opportunities to study how animals, in turn, shape ecosystems. We use high-spatiotemporal resolution Sentinel-2 satellite imagery to evaluate the effects of Arctic fox (*Vulpes lagopus*) denning activity on vegetation. Arctic fox dens are characterized with unique vegetation relative to the surrounding area, presumably due to decades of nutrient accumulation and bioturbation. We use an imagery-derived metric (NDVI) to compare maximum plant productivity and plant phenology patterns on Arctic fox dens vs. reference sites, i.e., points generated within areas of preferred denning habitat as predicted from a habitat selection analysis. We show that high-resolution satellite imagery can be used effectively to quantify the effects of Arctic fox denning activity on vegetation. Plant productivity and the rate of green up were both greater on fox dens compared to reference sites. Productivity on these preferred-habitat (reference) sites was lower than average productivity on the tundra (i.e., random sites), indicating that foxes primarily establish dens in low-productivity areas. Our findings support previous studies that proposed Arctic foxes function as ecosystem engineers in low Arctic ecosystems by converting sites of low productivity into sites of high productivity through their denning activity. Plant productivity was unrelated to recent den occupancy patterns, indicating fox denning activity has long-term legacy effects on plants that last well beyond the lifetime of foxes. We add to the growing body of literature that recognizes predators can be drivers of landscape heterogeneity and influence ecosystem dynamics through patch-scale pathways, such as by concentrating nutrients into localized areas. Our study demonstrates the efficacy of using remote sensing technologies to advance our understanding of the functional roles that predators specifically, and animals generally, occupy in ecosystems.

## Introduction

Technological and methodological advancements in remote sensing in recent decades have provided novel insights into relationships between animals and their environment. For instance, researchers can use airborne and satellite-based light detecting and ranging (LiDAR) data to remotely quantify and describe the influence of ecosystem 3D structure on animals (Davies & Asner 2014), including recent insights into habitat preferences of a threatened butterfly (de Vries et al. 2021) and how landscape heterogeneity mediates the competition and coexistence of sympatric predators (Davies et al. 2021). Satellite imagery is similarly used to evaluate the influence of environmental factors like precipitation, temperature, and land cover type on animals (Pettorelli et al. 2014). In particular, satellite-derived metrics such as the normalized difference vegetation index (NDVI) – a metric frequently used to approximate vegetation and ecosystem productivity – have been used to assess animal-habitat relationships across space and time (Pettorelli et al. 2005; Pettorelli et al. 2011). Most studies using satellite-derived metrics to advance animal ecology have been conducted on large spatial scales, including: a global-scale analysis on the influence of ecosystem productivity on scavenger assemblages (Sebastián-González et al. 2020); revealing how herbivores track vegetation green-up throughout migration corridors (Sawyer & Kauffman 2011; Bischof et al. 2012; van Moorter et al. 2013; Merkle et al. 2016; Aikens et al. 2017); how ecosystem productivity drives the structure of herbivore communities across the Arctic tundra (Speed et al. 2019); climate-driven effects on bird migration phenology (Saino et al. 2004; Gordo 2007); and modeling the distribution, abundance, and richness of species from numerous taxa (e.g., Hurlbert & Haskell 2003; Tognelli & Kelt 2004; Bartoń & Zalewski 2007; Evans et al. 2008; Nieto et al. 2015).

More recently launched satellites capable of capturing imagery at higher resolutions, including the Sentinel series launched by the European Space Agency, offer the potential to decipher animal-habitat relationships at finer spatial scales (Pettorelli et al. 2014). Compared to 30-m resolution data from current Landsat satellites, 10-m resolution imagery captured by the Sentinel-2 satellite was a better predicter of bird richness patterns across the continental United States (Farwell et al. 2021). Integrating these high-resolution data sources can also improve the performance of species distribution models (Koma et al. 2022) and help identify microhabitat selection and suitability for small animals (Valerio et al. 2020; Alessandrini et al. 2022). Many remote sensing data sources are also freely available and easily accessible via platforms like Google Earth Engine. With advancements in the technology, availability, and accessibility of these data, we can expect researchers, conservationists, and managers alike to use them to address old, new, and ever-evolving ecological questions (Turner et al. 2015; Schulte to Bühne & Pettorelli 2018). In particular, finer-scale data sources may help unravel not only how environmental conditions affect animals, but how animals, in turn, influence ecosystem dynamics.

Predators are widely recognized for their ecological influence on prey abundance and behavior but they also alter ecosystems through localized, patch-scale pathways that drive landscape heterogeneity across space and time. These pathways include distributing carcasses across the landscape (Bump et al. 2009; Schmitz et al. 2010; Risch et al. 2020; Monk & Schmitz 2022) and killing ecosystem engineers that create landscape patches (e.g., beaver [*Castor canadensis*] ponds; Gable et al. 2020), as well as concentrating nutrients derived from prey into discrete locations such as foraging (Holtgrieve et al. 2009), scent-marking (Ben-David et al. 1998; Crait & Ben-David 2007), and social aggregation sites (Fariña et al. 2003; Bokhorst et al. 2019). Through each of these pathways, predators alter or create patches that influence landscape heterogeneity by indirectly affecting other species in a localized manner. For instance, predators that perennially re-use home sites may indirectly affect local plant and soil communities by concentrating prey-derived nutrients there. Soil nutrient levels and plant growth are greater at ground-nesting eagle owl (*Bubo bubo*) nests compared to reference sites (Fedriani et al. 2015), whereas the combination of bioturbation (from digging burrows) and nutrient deposition by badgers (*Meles meles*) and red foxes (*Vulpes vulpes*) benefits plants around their dens (Kurek et al. 2014; Kucheravy et al. 2021; Lang et al. 2021). To date, few studies have looked at these patch-scale predator effects from a landscape-scale perspective (but see Bump et al. 2009; Gable et al. 2020).

Arctic foxes (*Vulpes lagopus*) are important terrestrial predators that occupy multiple functional roles in low Arctic tundra ecosystems. Although often recognized for their role in regulating the abundance (Angerbjörn et al. 1999; Bêty et al. 2001; Iles et al. 2013) and altering the behavior (Bêty et al. 2002; Clermont et al. 2021) of their prey, their distinctive dens have also garnered considerable scientific and public interest. Throughout parts of their range, Arctic fox dens are characterized by unique vegetation compared to the surrounding area (Fig. 1) (Chesemore 1969), earning them the nickname of the ‘gardens of the tundra’. Arctic fox dens have greater soil and plant nutrient content, greater plant biomass, and unique plant assemblages compared to nearby areas (Garrott et al. 1983; Smith et al. 1992; Bruun et al. 2005; Gharajehdaghipour et al. 2016; Gharajehdaghipour & Roth 2018; Fafard et al. 2020), and the foxes are likely the main drivers of these traits. Arctic foxes re-use dens for decades or even centuries (Macpherson 1969), and through time the combination of digging burrows and the accumulation of nutrients from fox excrement and prey remains likely alters local vegetation. As noted earlier, similar nutrient-enhancement patterns are found on other predator home sites (Kurek et al. 2014; Fedriani et al. 2015; Kucheravy et al. 2021; Lang et al. 2021), supporting the hypothesis that the predators are the cause of the enhanced vegetation. Arctic foxes, and other predators, have accordingly been classified as ecosystem engineers – organisms that benefit other species through physical modifications of their environment (Jones et al. 1994) – due to the unique vegetative traits localized to their dens. However, without accounting for den selection preferences at the landscape scale, it is theoretically possible that the predators chose high-productivity areas to build their dens.

**Figure 1.**
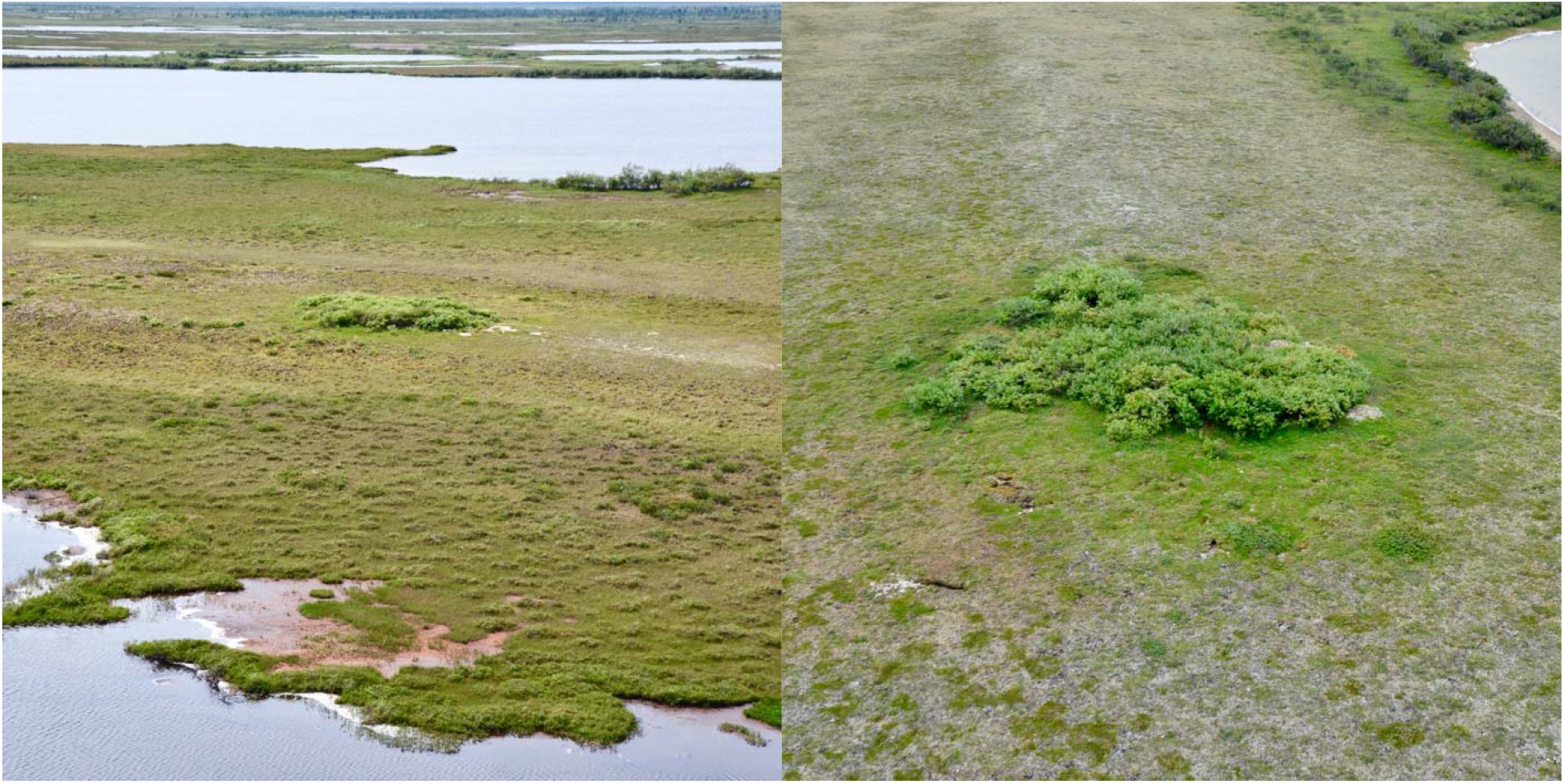
Photos of typical Arctic fox dens in Wapusk National Park, Manitoba, Canada taken during an aerial survey. Dens are generally constructed on relatively elevated ancient beach ridges and are characterized by unique plant characteristics that appear distinct against the surrounding landscape.

Here, we test the efficacy of using freely available, high spatial and temporal resolution remote sensing data to evaluate the effects of Arctic fox denning activity on vegetation, and ultimately unravel the role of Arctic foxes as ecosystem engineers in low Arctic tundra ecosystems. We first developed a habitat selection model for fox dens to create *reference* sites, which effectively represent areas suitable for denning based on factors influencing current fox den locations (i.e., preferred-habitat sites). Using NDVI data derived from Sentinel-2 satellite imagery (10 m resolution), we then compared 1) maximum plant productivity and 2) plant phenology patterns on (i) Arctic fox dens compared to (ii) reference sites on the tundra. We also compared plant productivity and phenology on fox dens and reference sites with (iii) fully random sites on the tundra (i.e., sites that represent all terrestrial habitats) to provide insight into how vegetation patterns on den and reference sites compare with average (random) tundra sites. We predicted plant productivity would be greater and plants would green up earlier and faster on fox dens compared to reference sites, supporting the hypothesis that Arctic foxes act as ecosystem engineers by altering local vegetation. Our study demonstrates the potential of using high-resolution remote sensing data to advance our understanding of the functional role(s) of predators in ecosystems.

## Methods

### Study Area

We conducted our study within a ∼1200 km^2^ tundra region of Wapusk National Park in northeastern Manitoba, Canada along the western coastline of Hudson Bay (Fig. 2A). Wapusk is located just east of the town of Churchill, Manitoba and is part of the Hudson Bay Lowlands, one of the largest wetland ecosystems in the world. Monthly average temperature ranges from - 26.0°C in January to 12.7°C in July (ECCC 2022). There is an average of 87 frost-free days (Jun. 19 to Sept. 15) annually in the area (ECCC 2022).

**Figure 2.**
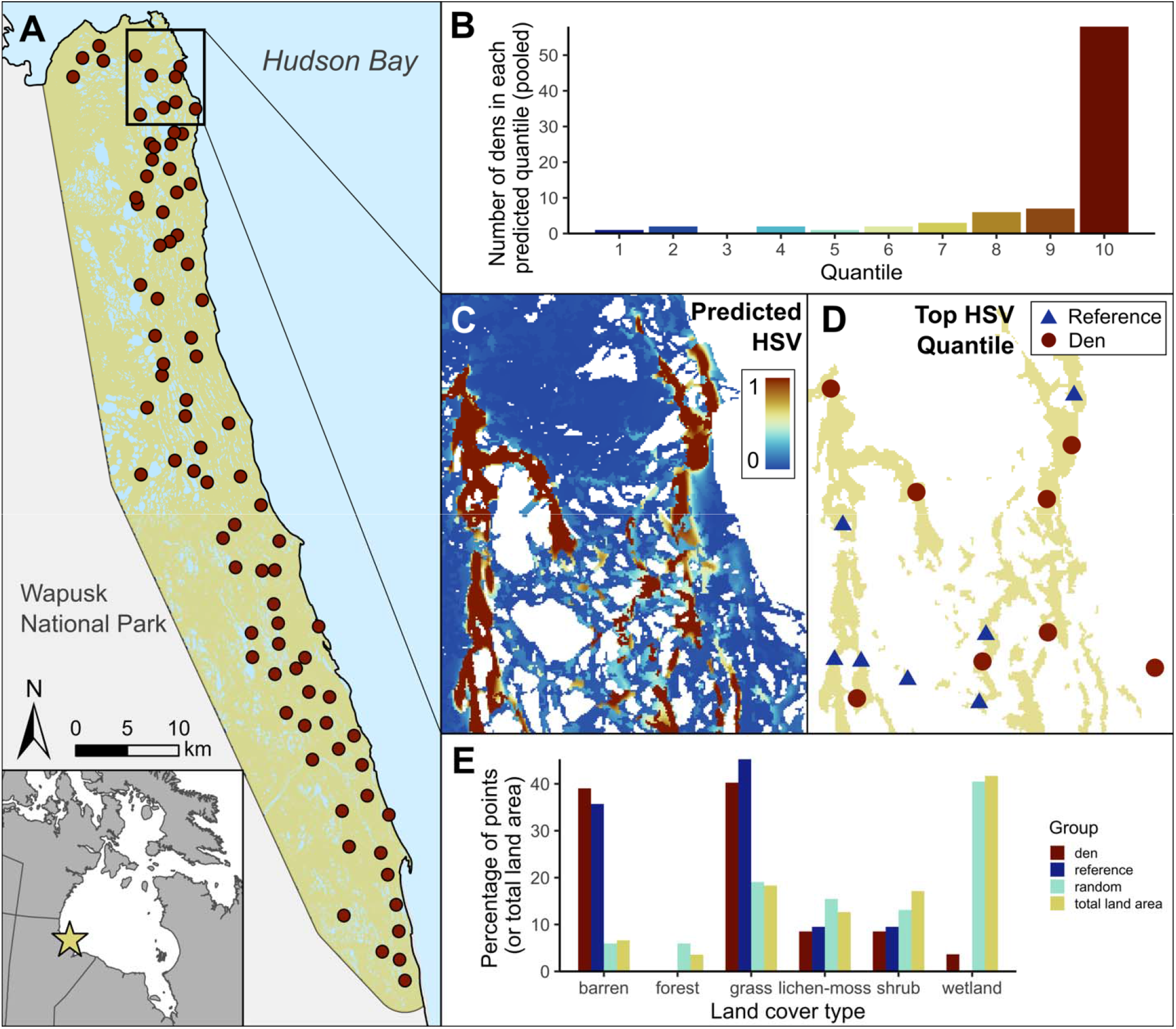
Map of the study area in Wapusk National Park, Manitoba, Canada, and results related to our den habitat selection analysis. Panel A shows the study area and all 84 known Arctic fox den locations therein. Panel B shows the number of Arctic fox dens predicted to be within each quantile (1–10) pooled across all five folds of the cross-validation procedure; Spearmans’s rank correlation (*r*) on the pooled quantiles was 0.86 (*p*=0.002). Panel C shows the predicted habitat selection map generated from the den selection model fit to the full data set, where habitat selection values (HSV) closer to 1 (red) represent areas more likely to be selected by Arctic foxes for denning. Panel D shows the same region with only the reclassified top quantile (10%) of cells present, overlaid with den (red circles) and randomly generated reference sites (blue triangles) used in this study. The study area region shown in Panels C and D is depicted within Panel A. Panel E shows the percentage of den, reference, and random points categorized by each of the six land cover types, along with the percentage of the total study area comprised of each cover type (not pictured: percent ‘open water’, which comprises ∼15% of the total study area).

Arctic fox dens in Wapusk are predominantly located on ancient beach ridges formed from isostatic rebound after the melting of the Keewatin ice sheet (Ritchie 1956; Roth 2003; Sella et al. 2007). The ridges are relatively elevated and run roughly parallel to the Hudson Bay shoreline, with ponds, lakes, and wetland habitats between the ridges. Beach ridges are thought to be suitable denning habitats for foxes due to the low soil moisture levels and greater depth to permafrost layer, which allows for easier burrow digging (Chesemore 1969; Smits et al. 1988; Dalerum et al. 2002). Our study area included 84 known Arctic fox dens that were used for the NDVI analysis (density: ∼7 dens/100 km^2^), but in the habitat selection analysis we excluded two dens that were misclassified as ‘open water’ according to the 2015 Canada Land Cover data set.

### Arctic fox den habitat selection analysis

To evaluate how Arctic foxes select denning locations, we first delineated the area available for possible denning sites by creating a minimum convex polygon around all known fox dens and then applying a 3-km buffer (clipped to the shoreline; Fig. 2A). This approach restricted the habitat selection analysis to only areas near known dens. Next, we generated 8200 random points (100 per den) within terrestrial habitats in the study area after removing all areas identified as ‘open water’ from the 2015 Canada Land Cover data set (Natural Resources Canada 2019).

We performed the habitat selection analysis by comparing den and random sites (input as 1 and 0, respectively) using a binomial generalized additive model with a logit link function. We used three variables for the analysis: elevation, land cover type, and latitude/longitude, the latter of which was used to help control for spatial autocorrelation. Elevation data was obtained from the 30-m resolution FABDEM data set (Hawker et al. 2022). We reclassified the 2015 Land Cover data (Natural Resources Canada 2019) into six categories: ‘forest’ (comprised of all forest cover types); ‘shrub’ (comprised of polar and non-polar shrub cover types); ‘grass’ (comprised of polar and non-polar grass cover types); ‘lichen-moss’; ‘wetland’; and ‘barren’. We used ‘wetland’ as the reference level, as it is the most abundant habitat type in the study area. The habitat selection model is defined as:

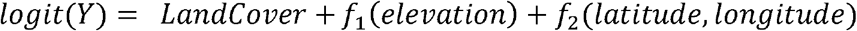

where *LandCover* is the point’s land cover type (categorical variable), *f*_*1*_*(elevation)* is the point’s elevation (in meters) fit with a smoothing component *f*_*1*_ using a cubic regression spline, and *f*_*2*_(*latitude, longitude)* is the interaction between the point’s latitude and longitude (in UTM units) fit with a smoothing component *f*_*2*_ using a Gaussian process spline.

#### Habitat selection model validation

Creating ‘reference’ points for the NDVI analysis relied upon having a habitat selection model that could adequately predict known Arctic fox den locations, as this model would be used to generate new points in locations where foxes are likely to create dens but have not done so yet. We therefore validated the habitat selection model performance using 5-fold cross-validation (detailed in Boyce et al. 2002; Roberts et al. 2017). Briefly, this process involved: 1) fitting 80% of the data to the habitat selection model, 2) creating a predictive habitat selection map (30-m resolution) of the study area from the model, 3) binning the map into 10 equal-sized quantiles, 4) identifying which quantile each den from the withheld (testing) data set was predicted to be in, and 5) performing a Spearman’s rank correlation test (R version 4.2.0; R Core Team 2022) on the withheld dens and their predicted quantile score. This process was repeated four more times until each 20% of data was withheld as a testing fold. We also performed a Spearman’s rank correlation on the predicted quantile data pooled across all five folds (Fig. 2B).

After model validation, which determined the habitat selection model performed adequately in predicting Arctic fox denning locations, we fit the full data set to the habitat selection model. We then created a predictive habitat selection map from this model (Fig. 2C), reclassified the map into 10 equal-sized quantiles, and generated 84 random points within cells of the top quantile >250m from another reference site and from the nearest den (Fig. 2D). These ‘reference’ points effectively represent the 10% of terrestrial areas Arctic foxes are most likely to select for denning. All habitat selection models were fit using the ‘gam’ function from the *mgcv* package (Wood 2011) in R.

### Plant productivity and phenology analyses

#### Evaluating maximum plant productivity

We compared maximum plant productivity between 84 each of 1) ‘den’ locations, 2) ‘reference’ locations representing preferred but unused denning habitat (described earlier), and 3) ‘random’ locations, which represent the total terrestrial habitat availability in the study area (Fig. 2E). ‘Random’ points were randomly generated in ArcGIS Pro (version 2.8; Esri 2022) within terrestrial habitats in the study area >250m from every other point (den, reference, and random), a distance that would ensure independence among the points.

We assessed maximum plant productivity by creating a greenest pixel mosaic of 2A surface reflectance Sentinel-2 imagery (captured every 2-4 days in our study area) across the full growing season for 2019, 2020, and 2021 using Google Earth Engine (2019 is the earliest year 2A imagery is available for our study area). We defined the growing season as Jun. 16–Sept. 30, which was based on the average dates of last and first frost (Jun. 19 and Sept. 15) and suitable satellite image availability (cloud and snow coverage interference is high outside of this date range). This process involved first extracting all satellite images within the growing season, applying the ‘s2cloudless’ (Zupanc 2017) algorithm to mask clouds and shadows from each image, calculating NDVI values for each image pixel, and then using the ‘qualityMosaic’ tool in Google Earth Engine to create a greenest pixel mosaic raster. This mosaic raster represents the maximum NDVI value on a pixel-by-pixel basis across the growing season. We exported the mosaic raster to ArcGIS Pro, excluded all cells with NDVI values <0.1 (as these cells corresponded to open water features or noise in the data), and calculated the mean NDVI value within a 20m buffer of each point (example in Fig. 3C) using the ‘Zonal Statistics as Table’ tool. We compared 20m maximum NDVI values between dens, reference points, and random points using a linear mixed effects model with ‘point ID’ and ‘year’ as random intercept terms (*lme4* package; Bates et al. 2015). Tukey’s Honest Significant Difference pairwise comparisons were conducted using the ‘pairs’ function from the *emmeans* package (Lenth 2022). Notably, the average den size in our study area is ∼563 m^2^, so the 20-m radius (1257 m^2^) includes area beyond many dens; NDVI estimates for dens should therefore be conservative.

**Figure 3.**
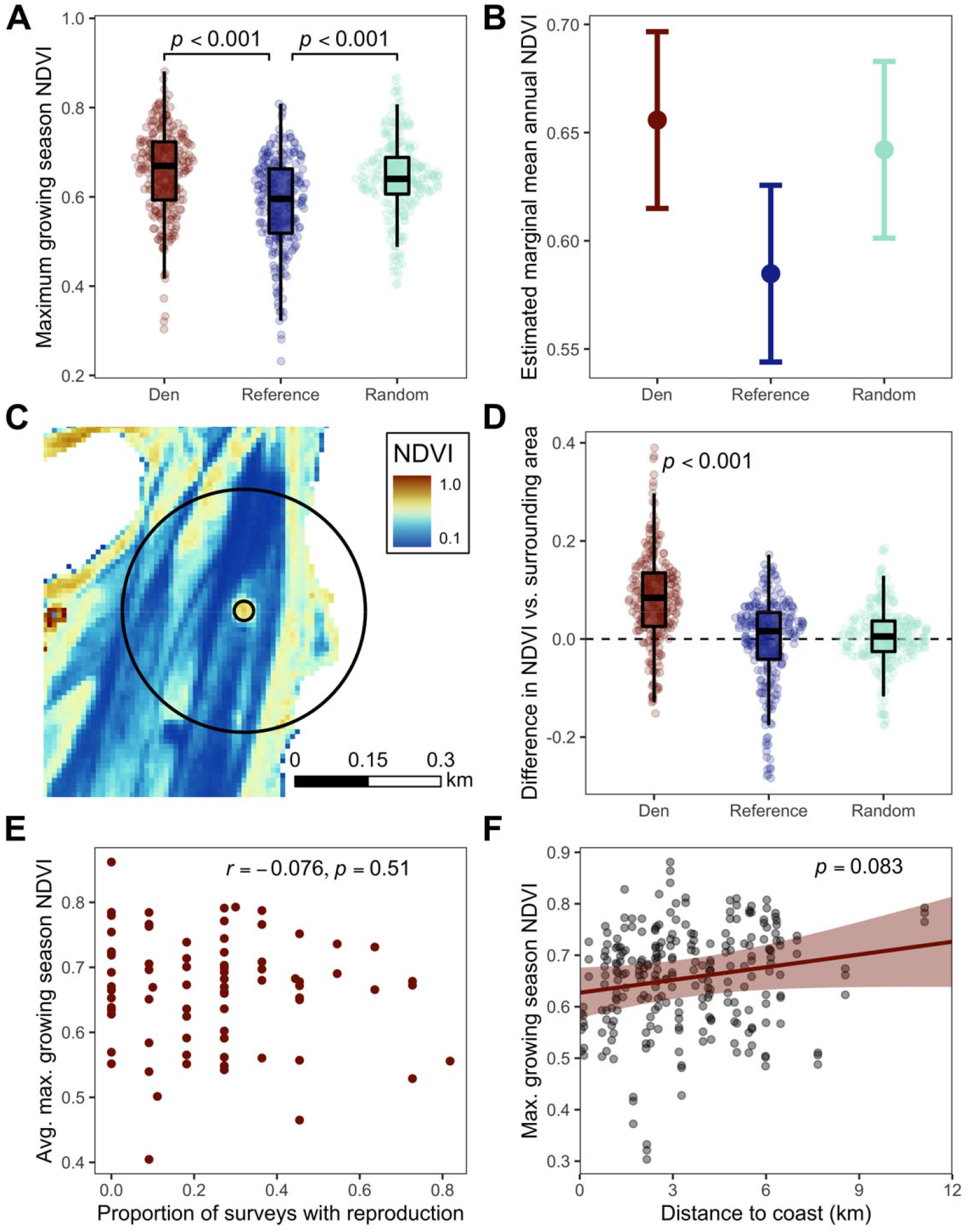
Results related to maximum plant productivity analyses on Arctic fox dens in Wapusk National Park, Manitoba, Canada. Panels A and B show the observed values (A) and estimated marginal mean (+/-95% confidence intervals; B) of maximum growing season NDVI for dens, reference sites, and random sites, where reference sites represent preferred denning habitats based on a habitat selection model and random sites are representative of total habitat availability in the study area. Panel C shows how NDVI scores were calculated within 20 and 250 m buffers (white portions are open water areas that were excluded from all analyses). The buffers are centered around an Arctic fox den. Panel D shows the observed difference between annual average NDVI values within 20 and 250 m buffers, with differences only found for dens. Panel E shows the lack of relationship between fox reproductive success and average maximum growing season NDVI on dens, while F shows the weak relationship between distance to coast and maximum NDVI.

We were also interested in quantifying how prominent Arctic fox dens are on the landscape relative to the other locations used in the productivity analysis (reference and random groups). We therefore compared the difference in mean NDVI values within a 20m buffer vs. mean NDVI values within a 250m buffer for each point, each year (2019-2021). We averaged the mean maximum NDVI for each point across the 3 years and used Wilcoxon signed rank tests in R to assess differences in average productivity between the two buffer distances for each group.

Finally, we assessed how recent fox reproductive success and each den’s distance to the coast affected maximum plant productivity on dens. For each year spanning 2011–2021, we assessed the reproductive success of dens using on-the-ground and aerial surveys (details in McDonald et al. 2017). Briefly, we examined each den for signs of reproductive success, such as abundant prey remains on dens, fresh digging in burrows, and presence of fresh pup scats. To evaluate whether recent fox reproduction patterns influenced plant productivity, we used a Spearman’s rank correlation test to evaluate the relationship between percent reproductive success (defined as the number of years each den produced pups divided by the number of years the den was surveyed) and maximum growing season NDVI averaged over the three years of imagery. We included only dens that had been surveyed at least 9 times for this analysis (*n*=78 dens). A previous study from this region demonstrated shrub occurrence is greater on dens farther from the coast (Fafard et al. 2020), so we hypothesized productivity may likewise be greater on dens farther inland. We evaluated the influence of distance to coast on 20m buffer NDVI values for all dens using a linear mixed effects model with ‘den ID’ and ‘year’ as random intercept terms.

#### Evaluating patterns of plant phenology

To understand whether plant phenology patterns on Arctic fox dens differed from other areas on the tundra, we examined whether dens green-up earlier or stay green longer, and when, during the growing season, any differences in plant productivity may arise. For this analysis, we used a similar ‘greenest pixel mosaic’ approach as described for the plant productivity analyses. We divided the growing season into seven equal time periods (Jun. 16-30, Jul. 1-15, Jul. 16-31, Aug. 1-15, Aug. 16-31, Sep. 1-15, Sep. 16-30) and generated greenest pixel NDVI mosaics for each time period spanning 2019-2021. Because of issues with cloud and shadow interference in the mosaics generated from the shorter time periods of this analysis, we manually inspected each mosaic raster and retained only the points where NDVI values could be satisfactorily calculated. For each time period raster mosaic, we calculated mean NDVI within a 20m buffer of each point using the methods described earlier. We evaluated plant NDVI phenology using a hierarchical generalized additive model with a Gaussian distribution:

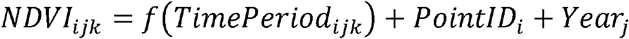

where *NDVI*_*ijk*_ is the mean 20m NDVI value of the *k*th observation for point *i* in year *j*, and *PointID* and *Year* are random intercept terms that were assumed to be normally distributed with a mean of 0. *f* (*TimePeriod*_*ijk*_)is the time period of the *k*th observation for point *i* in year *j* (coded as an integer from 1 to 7, representing the seven equal time periods) fit with a smoothing component *f* using a thin plate regression spline comprised of seven basis functions. The spline varied by group level (den, reference, random) and had individual penalties (i.e., group-level smoothers with no shared penalty; Pedersen et al. 2019). The phenology model was fit using the ‘gam’ function in the *mgcv* package (Wood 2011).

## Results

### Fox den habitat selection

Arctic foxes predominantly constructed dens in ‘barren’ or ‘grass’ land cover types and avoided ‘wetland’ areas (Fig. 2E). The smoothing components of elevation (effective degrees of freedom [edf]=4.90, Chi-square=66.4, *p*<0.001) and the interaction between latitude/longitude parameters (edf=18.79, Chi-square=84.4, *p*<0.001) were both significant, with fox dens occurring more frequently in elevated areas nearer to the coast than random sites. The influence of proximity to the coastline on den selection was likely due in part to how the study area was delineated, which included inland areas west of known dens and removed areas in the bay that would otherwise have been included within the 3km buffer (Fig. 2A).

#### Model validation results

Results from our five-fold cross-validation procedure indicated the den habitat selection model adequately predicts Arctic fox den locations. Spearman’s rank correlation (*r*) results ranged from 0.43 (*p*=0.22) to 0.81 (*p*=0.005), with an average *r* of 0.67. However, the variation in *r* values may have been due in large part to the low sample size of dens used for each testing fold (*n*=16 or 17), whereby 1-2 dens predicted into low quantiles can greatly influnce *r* values. When pooled across all five folds, the Spearman’s rank correlation test on the predicted quantiles was significant (*r*=0.86, *p*=0.002), with 71% (58/82) and 87% (71/82) of dens predicted to be within the top one and three quantiles, respectively (Fig. 2B). When fit to the full data set the habitat selection model explained 31.8% of deviance in Arctic fox den charactersistics. We interpreted these results to indicate the habitat selection model had a moderate to good ability to explain current fox den characteristics and could be used to adequately predict other areas foxes may select for when constructing new dens (i.e., reference sites). The similarity in land cover types between den and reference sites within the top quantile supports this interpretation (Fig. 2E).

### Maximum plant productivity

Maximum growing season plant productivity was significantly greater on Arctic fox dens compared to reference sites (*p*<0.001, *T*=4.89; Fig. 3A). The estimated marginal mean annual NDVI was 0.66 (95% confidence interval [CI]: 0.62–0.70) on dens compared to 0.59 (95% CI: 0.54–0.63) for reference sites (Fig. 3B). Plant productivity at random sites was significantly greater than productivity at reference sites (*p*<0.001, *T*=-3.94; Fig. 3A-B), suggesting that Arctic fox dens are typically constructed in relatively low-productivity areas. There was no difference in plant productivity between den and random sites (*p*=0.61, *T*=0.94).

Plant productivity was significantly greater on Arctic fox dens compared to the surrounding area, such that average maximum NDVI values within a 20m buffer around Arctic fox dens were greater than average NDVI values within a 250m buffer (*T*=8.07, *p*<0.001, df=83; Fig. 3D). As expected, plant productivity at reference and random sites did not differ from their surrounding areas (*p*=0.58 and 0.33, respectively; Fig. 3D).

Average fox den reproductive success (number of years pups were produced at each den divided by the number of years surveyed) over the last 11 years was 0.25 (SD = 0.20). We found no relationship between recent fox reproduction and plant productivity on dens (Spearman’s *r*=- 0.076, *p*=0.51; Fig. 3E). There was also weak evidence for greater plant productivity on dens farther from the coastline (*Z*=1.734, *p*=0.083, marginal *R*^*2*^=0.031; Fig. 3F).

### Plant phenology

We found Sentinel-2 imagery can be used to detect and visualize plant phenology patterns on Arctic fox dens (Fig. 4A). Cloud and snow coverage regularly interfered with obtaining data from individual points, but across all seven time periods spanning 2019–2021 we acquired 3,779 useable NDVI data points (71.4% total success rate in acquiring data to calculate NDVI, with a range across the seven time periods of 23.8% to 97.2%).

**Figure 4.**
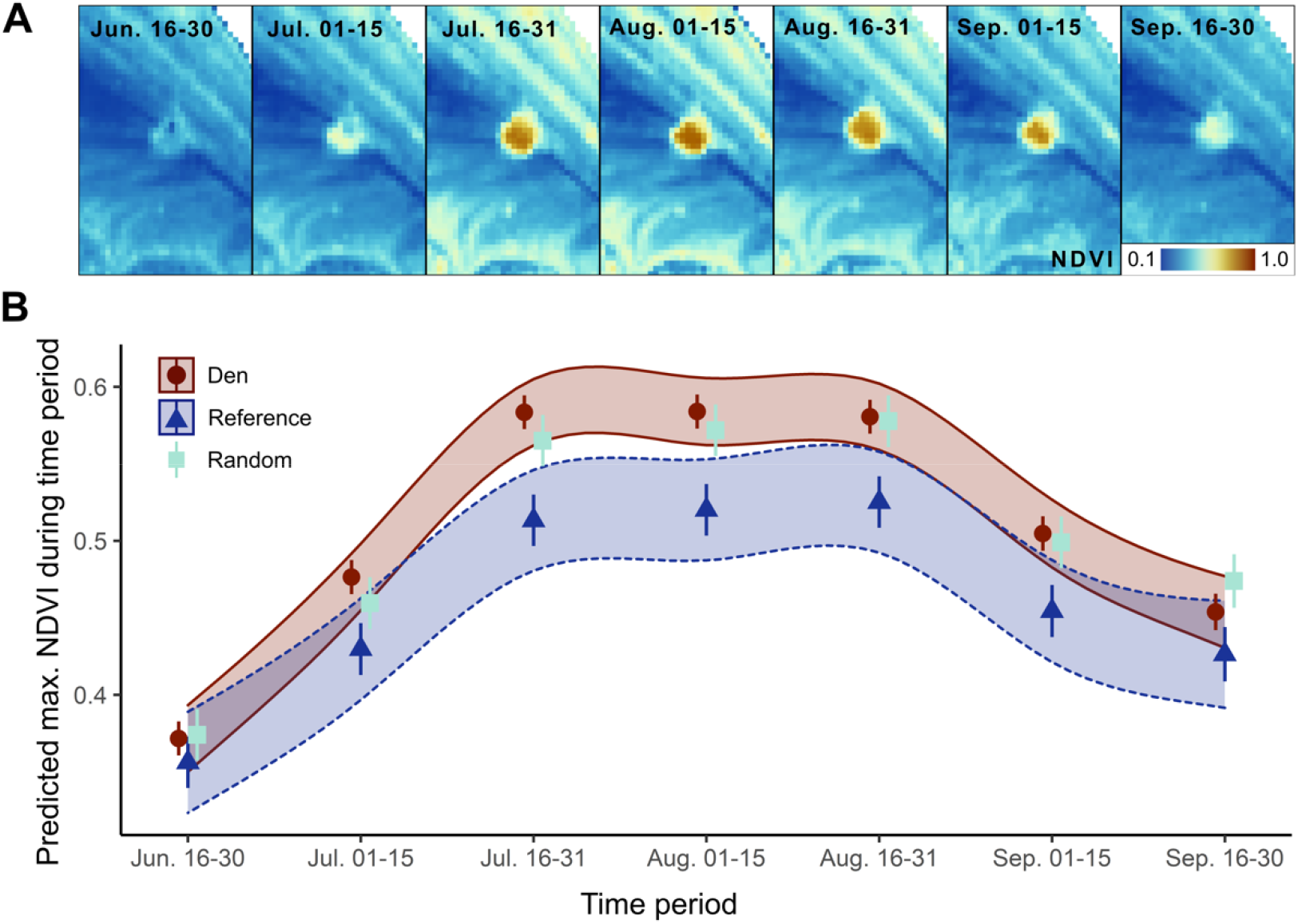
Results related to the plant phenology analysis on Arctic fox dens in Wapusk National Park, Manitoba, Canada. Panel A shows the intra-annual change in plant productivity (assessed from NDVI values) across seven time periods on a single Arctic fox den. Each subpanel shows the NDVI values created from the greenest pixel mosaic from each time period. Panel B shows the predicted point and standard error NDVI estimates for den, reference, and random sites (generated using the *marginaleffects* R package; Arel-Bundock 2022). The ribbons are the predicted NDVI 95% confidence intervals (CI) for den and reference sites (the 95% CI ribbon for random sites was similar to the den ribbon and not shown here for easier interpretation).

As expected, the temporal trends in plant productivity (NDVI) across the growing season were all statistically significant and non-linear for each group (den, reference, and random sites). Specifically, plant productivity increased from mid-June until a peak around mid-July, where it remained at peak productivity until the end of August (Fig. 4B). Differences in plant productivity between den and reference sites were greatest during peak growing season (Fig. 4B), matching results from the maximum productivity analyses. We found no evidence that plants green-up earlier or stay green longer on dens relative to reference sites; however, the rate of plant green-up appears to be greater on dens (Fig. 4B).

## Discussion

Using freely available software and high-resolution satellite imagery, our study provides a novel, landscape-scale perspective on the effects of Arctic fox denning activity on plant productivity and phenology. We demonstrated that plant productivity on Arctic fox dens is significantly greater than other areas in similar habitats, which are generally limited to relatively elevated but low-productivity areas. By using a habitat selection analysis to generate reference points, we were able to control for certain ecological factors that Arctic foxes select for when creating dens and thus help disentangle the relative effects of habitat vs. fox denning activity on plants. Our results provide further evidence that Arctic foxes are ecosystem engineers in low Arctic tundra ecosystems.

### Which came first: the foxes or the plants?

One of the lingering questions related to vegetation patterns on Arctic fox dens is whether their denning activity converts unproductive sites into productive sites through nutrient deposition and bioturbation, or whether they select for preexisting high-productivity sites to build their dens. Despite previous findings of unique plant characteristics on Arctic fox dens (Smith et al. 1992; Bruun et al. 2005; Gharajehdaghipour et al. 2016; Gharajehdaghipour & Roth 2018; Fafard et al. 2020), these analyses were conducted on smaller spatial scales that compared these metrics with immediately adjacent areas and thus could not rule out whether foxes selected locations with relatively high plant productivity. By generating reference points based on an Arctic fox den habitat selection model that controlled for factors influencing den site characteristics, we show that Arctic foxes select for low-productivity areas when digging dens: plant productivity at reference points was significantly less than at random points on the tundra (Fig. 3A-B). Arctic foxes could still select for relatively productive spots within unproductive habitats, but in that case we should have detected little difference in plant productivity between dens and reference sites. Although we still cannot say with total certainty that the foxes precede the vegetation change at den sites, the fact that plant productivity was greater on Arctic fox dens than reference sites provides further evidence that Arctic foxes are responsible for the unique plant patterns on dens, as suggested previously (Gharajehdaghipour et al. 2016; Gharajehdaghipour & Roth 2018; Fafard et al. 2020). Our landscape-scale analysis supports the notion that Arctic foxes act as ecosystem engineers by converting sites of low productivity into sites of relatively high productivity through their denning activity.

The unique plant characteristics on Arctic fox dens makes them visually prominent patches on the elevated beach ridges, which are otherwise largely unvegetated or comprised of prostrate shrubs. Our analysis supports what the ‘eye test’ suggests: plant productivity is significantly greater on Arctic fox dens compared to the immediately surrounding area (Fig. 3D), which drives how visually distinct these engineered patches are. As expected, we did not see the same pattern in reference or random sites; our sampling design was conducted on a random and semi-random basis rather than deliberately choosing relatively high productive patches. Our objective with this portion of the analysis was to derive some index of ‘prominence’ that could quantify what our eyes perceive when viewing these dens.

The fact that fox reproduction patterns spanning a decade are unrelated to maximum plant productivity on dens indicates the effects of fox denning behavior on plants are long-lasting – well beyond the lifetime of foxes. Indeed, the effects from nutrient deposition and bioturbation seem to compound through many generations of fox occupancy and reproduction. And once the changes to plant assemblages occur on dens, these legacy effects likely last for a long time. Continued monitoring of plant productivity and assemblages may reveal greater information about how plant dynamics change through time on Arctic fox dens in relation to occupancy patterns (see *Future directions and concluding remarks* section for more discussion).

### Harnessing high-resolution remote sensing data to assess animal-habitat relationships

Our first objective when planning this study was to conduct a “proof of concept” analysis evaluating whether satellite imagery could actually detect the unique vegetation on Arctic fox dens. Prior to the launch of the Sentinel-2 satellite, freely available, high-temporal frequency imagery was available only on a coarser scale (e.g., 30-m resolution of the LANDSAT satellites). Arctic fox dens are represented by ∼1 pixel at this resolution (900 m^2^), making it challenging to detect dens at all let alone spatiotemporal vegetation patterns. Sentinel-2 imagery clearly provides a fine enough spatiotemporal scale to detect and quantify plant productivity and phenology patterns on Arctic fox dens (Figs. 3, 4). Although plants on dens do not appear to green-up earlier or stay green longer than other tundra areas, we found green-up rates were higher on dens than reference sites and were able to assess when, during the growing season, plant productivity begins to differ between these areas (∼mid-July, near the peak growing season; Fig. 4B). Assessing plant phenology patterns on these dens at a similar spatiotemporal scale using field-based methods would have been prohibitively costly and time intensive, as the majority of the dens in our 1200-km^2^ study area (∼65-70%) are accessible only via helicopter during the growing season.

As remote sensing data further increases in availability and spatiotemporal resolution, it will continue to offer new perspectives and insights on animal functional roles within ecosystems. Most animal ecology studies using remote sensing data have applied it towards understanding how environmental conditions affect animals. But as we and others have demonstrated, remote sensing data can also be used to understand how animals shape ecosystem dynamics themselves. For instance, satellite imagery has been used to identify how spatiotemporal fluctuations in spawning salmon abundance influences forest productivity (Kieran et al. 2021), how beaver-modified environments buffer riparian ecosystems against wildfire (Fairfax & Whittle 2020), and how grazing pressure from migrating bison (*Bison bison*) alters the quality and phenology of the grasses they forage upon (Geremia et al. 2019). Airborne LiDAR has revealed the effects of elephant (*Loxodonta africana*) foraging on the structural diversity, rates of treefall, and vegetation height of savanna woodlands (Asner et al. 2009; Asner & Levick 2012; Asner et al. 2016; Davies et al. 2018). We add to these studies by demonstrating high-resolution satellite imagery can provide a landscape-scale perspective of animal ecological effects that function at smaller, patch-level scales.

### Future directions and concluding remarks

By demonstrating that satellite imagery can be used to detect vegetation patterns on Arctic fox dens, there are several ways this study can be built upon for future research and monitoring efforts. First, we propose satellite-derived information could be used to identify and locate previously unknown dens with similar vegetation characteristics. In particular, the prominence index we employed may be used to identify vegetation hotspots that, in tandem with region-specific den selection preferences, may guide search efforts for dens. Second, as we continue to assess fox occupancy and reproductive success at dens, we will likely gain a better understanding of how fox denning behavior affects plants across space and time. For instance, how long do the changes to plant communities last once a den is abandoned for good? After a new den is constructed, how long until major changes to plants are detectable? Finally, as Sentinel-2 and other comparable satellites continue to collect high-resolution spatiotemporal imagery, these data can be used to regularly monitor temporal patterns in plant productivity and phenology on dens and how climate change may affect these patterns across multiple spatial scales.

Effective species conservation and management often requires employing efficient ways to monitor and measure animal-habitat relationships. This is especially true when resources are limited, when studies are conducted across large spatial or temporal scales, or when the species are imperiled or have important ecological effects, like top predators do. Remote sensing data offer unique avenues to advance these objectives (Pettorelli et al. 2014; Schulte to Bühne & Pettorelli 2018), provided the data remain affordable and accessible (Turner et al. 2015). Our study validates the utility of such data by demonstrating how free and easily accessible satellite imagery can provide a cosmic view of predator ecological impacts, and ultimately advance our understanding of the intricate functional role predators play in ecosystems.

## Acknowledgments

We acknowledge salary and stipend support provided by the University of Manitoba. Support in locating and monitoring Arctic fox dens was provided by the Natural Sciences and Engineering Research Council of Canada, Natural Resources Canada Polar Continental Shelf Program, University of Manitoba Fieldwork Support Program, the Churchill Northern Studies Centre Northern Research Fund, and the many students that hiked out to dens as part of the Churchill Fox Project long-term research and monitoring efforts.

## Data availability

Due to the sensitive nature of Arctic fox dens, specific location data is only available upon request from the authors. Otherwise, all other data and code used in this study will be available from the Dryad Digital Repository.

## Author contributions

S.M. Johnson-Bice conceived the ideas, designed the methodology, collected the data, analyzed the data, and led the writing of the manuscript. J.D. Roth and J.H. Markham supported in conceiving the study ideas and designing the methodology. All authors contributed critically to the drafts and gave final approval for publication.

## References

Aikens, E.O. et al. (2017) The greenscape shapes surfing of resource waves in a large migratory herbivore. Ecology Letters, 20, 741–750.

Alessandrini, C., Scridel, D., Boitani, L., Pedrini, P. & Brambilla, M. (2022) Remotely sensed variables explain microhabitat selection and reveal buffering behaviours against warming in a climate-sensitive bird species. Remote Sensing in Ecology and Conservation, n/a.

Angerbjörn, A., Tannerfeldt, M. & Erlinge, S. (1999) Predator-prey relationships: arctic foxes and lemmings. Journal of Animal Ecology, 68, 34–49.

Arel-Bundock, V. (2022) marginaleffects: Marginal effects, marginal means, predictions, and contrasts. R package version 0.6.0. https://CRAN.R-project.org/package=marginaleffects.

Asner, G. P. et al. (2009) Large-scale impacts of herbivores on the structural diversity of African savannas. Proceedings of the National Academy of Sciences, 106, 4947–4952.

Asner, G.P. & Levick, S.R. (2012) Landscape-scale effects of herbivores on treefall in African savannas. Ecology Letters, 15, 1211–1217.

Asner, G.P., Vaughn, N., Smit, I.P.J. & Levick, S. (2016) Ecosystem-scale effects of megafauna in African savannas. Ecography, 39, 240–252.

Bartoń, K.A. & Zalewski, A. (2007) Winter severity limits red fox populations in Eurasia. Global Ecology and Biogeography, 16, 281–289.

Bates, D., Mächler, M., Bolker, B. & Walker, S. (2015) Fitting linear mixed-effects models using lme4. Journal of Statistical Software, 67, 1–48.

Ben-David, M., Bowyer, R.T., Duffy, L.K., Roby, D.D. & Schell, D.M. (1998) Social behavior and ecosystem processes: river otter latrines and nutrient dynamics of terrestrial vegetation. Ecology, 79, 2567–2571.

Bêty, J., Gauthier, G., Giroux, J.-F. & Korpimäki, E. (2001) Are goose nesting success and lemming cycles linked? Interplay between nest density and predators. Oikos, 93, 388–400.

Bêty, J., Gauthier, G., Korpimäki, E. & Giroux, J.-F. (2002) Shared predators and indirect trophic interactions: lemming cycles and arctic-nesting geese. Journal of Animal Ecology, 71, 88–98.

Bischof, R. et al. (2012) A Migratory Northern Ungulate in the Pursuit of Spring: Jumping or Surfing the Green Wave? The American Naturalist, 180, 407–424.

Bokhorst, S., Convey, P. & Aerts, R. (2019) Nitrogen inputs by marine vertebrates drive abundance and richness in Antarctic terrestrial ecosystems. Current Biology, 29, 1721-1727.e1723.

Boyce, M.S., Vernier, P.R., Nielsen, S.E. & Schmiegelow, F.K.A. (2002) Evaluating resource selection functions. Ecological Modelling, 157, 281–300.

Bruun, H.H., Österdahl, S., Moen, J. & Angerbjörn, A. (2005) Distinct patterns in alpine vegetation around dens of the Arctic fox. Ecography, 28, 81–87.

Bump, J.K., Peterson, R.O. & Vucetich, J.A. (2009) Wolves modulate soil nutrient heterogeneity and foliar nitrogen by configuring the distribution of ungulate carcasses. Ecology, 90, 3159–3167.

Chesemore, D.L. (1969) Den ecology of the Arctic fox in northern Alaska. Canadian Journal of Zoology, 47, 121–129.

Clermont, J. et al. (2021) The predator activity landscape predicts the anti-predator behavior and distribution of prey in a tundra community. Ecosphere, 12, e03858.

Crait, J.R. & Ben-David, M. (2007) Effects of river otter activity on terrestrial plants in trophically altered yellowstone lake. Ecology, 88, 1040–1052.

Dalerum, F., Tannerfeldt, M., Elmhagen, B., Becker, D. & Angerbjörn, A. (2002) Distribution, morphology and use of arctic fox Alopex lagopus dens in Sweden. Wildlife Biology, 8, 185–192.

Davies, A.B. & Asner, G.P. (2014) Advances in animal ecology from 3D-LiDAR ecosystem mapping. Trends in Ecology & Evolution, 29, 681–691.

Davies, A.B., Gaylard, A. & Asner, G.P. (2018) Megafaunal effects on vegetation structure throughout a densely wooded African landscape. Ecological Applications, 28, 398–408.

Davies, A.B. et al. (2021) Spatial heterogeneity facilitates carnivore coexistence. Ecology, 102, e03319.

de Vries, J.P.R., Koma, Z., WallisDeVries, M.F. & Kissling, W.D. (2021) Identifying fine-scale habitat preferences of threatened butterflies using airborne laser scanning. Diversity and Distributions, 27, 1251–1264.

Environment Climate Change Canada (ECCC) (2022) Climate Normals 1981-2010 for the Churchill, MB station. Accessed 4 Aug. 2022.

ESRI (2022) ArcGIS Pro version 2.8. Redlands, CA.

Evans, K.L., Newson, S.E., Storch, D., Greenwood, J.J.D. & Gaston, K.J. (2008) Spatial scale, abundance and the species–energy relationship in British birds. Journal of Animal Ecology, 77, 395–405.

Fafard, P.M., Roth, J.D. & Markham, J.H. (2020) Nutrient deposition on Arctic fox dens creates atypical tundra plant assemblages at the edge of the Arctic. Journal of Vegetation Science, 31, 173–179.

Fairfax, E. & Whittle, A. (2020) Smokey the Beaver: beaver-dammed riparian corridors stay green during wildfire throughout the western USA. Ecological Applications, 30, e02225.

Fariña, J.M., Salazar, S., Wallem, K.P., Witman, J.D. & Ellis, J.C. (2003) Nutrient exchanges between marine and terrestrial ecosystems: the case of the Galapagos sea lion Zalophus wollebaecki. Journal of Animal Ecology, 72, 873–887.

Farwell, L.S. et al. (2021) Satellite image texture captures vegetation heterogeneity and explains patterns of bird richness. Remote Sensing of Environment, 253, 112175.

Fedriani, J.M., Garrote, P.J., Delgado, M.d.M. & Penteriani, V. (2015) Subtle gardeners: inland predators enrich local topsoils and enhance plant growth. PLoS ONE, 10, e0138273.

Gable, T.D., Johnson-Bice, S.M., Homkes, A.T., Windels, S.K. & Bump, J.K. (2020) Outsized effect of predation: Wolves alter wetland creation and recolonization by killing ecosystem engineers. Science Advances, 6, eabc5439.

Garrott, R.A., Eberhardt, L.E. & Hanson, W.C. (1983) Arctic fox den identification and characteristics in northern Alaska. Canadian Journal of Zoology, 61, 423–426.

Geremia, C. et al. (2019) Migrating bison engineer the green wave. Proceedings of the National Academy of Sciences USA, 116, 25707–25713.

Gharajehdaghipour, T. & Roth, J.D. (2018) Predators attract prey through ecosystem engineering in the Arctic. Ecosphere, 9, e02077.

Gharajehdaghipour, T., Roth, J.D., Fafard, P.M. & Markham, J.H. (2016) Arctic foxes as ecosystem engineers: increased soil nutrients lead to increased plant productivity on fox dens. Scientific Reports, 6, 24020.

Gordo, O. (2007) Why are bird migration dates shifting? A review of weather and climate effects on avian migratory phenology. Climate Research, 35, 37–58.

Hawker, L. et al. (2022) A 30 m global map of elevation with forests and buildings removed. Environmental Research Letters, 17, 024016.

Holtgrieve, G.W., Schindler, D.E. & Jewett, P.K. (2009) Large predators and biogeochemical hotspots: brown bear (Ursus arctos) predation on salmon alters nitrogen cycling in riparian soils. Ecological Research, 24, 1125–1135.

Hurlbert, Allen H. & Haskell John P. (2003) The Effect of Energy and Seasonality on Avian Species Richness and Community Composition. The American Naturalist, 161, 83–97.

Iles, D.T. et al. (2013) Predators, alternative prey and climate influence annual breeding success of a long-lived sea duck. Journal of Animal Ecology, 82, 683–693.

Jones, C.G., Lawton, J.H. & Shachak, M. (1994) Organisms as ecosystem engineers. Oikos, 69, 373–386.

Kieran, C.N., Obrist, D.S., Muñoz, N.J., Hanly, P.J. & Reynolds, J.D. (2021) Links between fluctuations in sockeye salmon abundance and riparian forest productivity identified by remote sensing. Ecosphere, 12, e03699.

Koma, Z. et al. (2022) Better together? Assessing different remote sensing products for predicting habitat suitability of wetland birds. Diversity and Distributions, 28, 685–699.

Kucheravy, C.E., Roth, J.D. & Markham, J.H. (2021) Red foxes increase reproductive output of white spruce in a non-mast year. Basic and Applied Ecology, 51, 11–19.

Kurek, P., Kapusta, P. & Holeksa, J. (2014) Burrowing by badgers (Meles meles) and foxes (Vulpes vulpes) changes soil conditions and vegetation in a European temperate forest. Ecological Research, 29, 1–11.

Lang, J.A., Roth, J.D. & Markham, J.H. (2021) Foxes fertilize the subarctic forest and modify vegetation through denning. Scientific Reports, 11, 3031.

Lenth, R. (2022) emmeans: Estimated Marginal Means, aka Least-Squares Means. R package version 1.7.5. https://CRAN.R-project.org/package=emmeans.

Macpherson, A.H. (1969) The dynamics of Canadian arctic fox populations. Canadian Wildlife Service Report Series No. 8. Ottawa, CA. p52 pp.

McDonald, R.S., Roth, J.D. & Baldwin, F.B. (2017) Goose persistence in fall strongly influences Arctic fox diet, but not reproductive success, in the southern Arctic. Polar Research, 36, sup1.5.

Merkle, J.A. et al. (2016) Large herbivores surf waves of green-up during spring. Proceedings of the Royal Society B: Biological Sciences, 283, 20160456.

Monk, J.D. & Schmitz, O.J. (2022) Landscapes shaped from the top down: predicting cascading predator effects on spatial biogeochemistry. Oikos, 2022, e08554.

Natural Resources Canada (2019) 2015 Land Cover of Canada. Canada Centre for Remote Sensing, Natural Resources Canada, Government of Canada

Nieto, S., Flombaum, P. & Garbulsky, M.F. (2015) Can temporal and spatial NDVI predict regional bird-species richness? Global Ecology and Conservation, 3, 729–735.

Pedersen, E.J., Miller, D.L., Simpson, G.L. & Ross, N. (2019) Hierarchical generalized additive models in ecology: an introduction with mgcv. PeerJ, 7, e6876.

Pettorelli, N. et al. (2014) Satellite remote sensing for applied ecologists: opportunities and challenges. Journal of Applied Ecology, 51, 839–848.

Pettorelli, N. et al. (2011) The Normalized Difference Vegetation Index (NDVI): unforeseen successes in animal ecology. Climate Research, 46, 15–27.

Pettorelli, N. et al. (2005) Using the satellite-derived NDVI to assess ecological responses to environmental change. Trends in Ecology & Evolution, 20, 503–510.

R Core Team (2022) R: a language and environment for statistical computing [Version 4.2]. R Foundation for Statistical Computing, Vienna, Austria.

Risch, A.C. et al. (2020) Effects of elk and bison carcasses on soil microbial communities and ecosystem functions in Yellowstone, USA. Functional Ecology, 34, 1933–1944.

Ritchie, J.C. (1956) The native plants of Churchill, Manitoba, Canada. Canadian Journal of Botany, 34, 269–320.

Roberts, D.R. et al. (2017) Cross-validation strategies for data with temporal, spatial, hierarchical, or phylogenetic structure. Ecography, 40, 913–929.

Roth, J.D. (2003) Variability in marine resources affects arctic fox population dynamics. Journal of Animal Ecology, 72, 668–676.

Saino, N. et al. (2004) Ecological conditions during winter predict arrival date at the breeding quarters in a trans-Saharan migratory bird. Ecology Letters, 7, 21–25.

Sawyer, H. & Kauffman, M.J. (2011) Stopover ecology of a migratory ungulate. Journal of Animal Ecology, 80, 1078–1087.

Schmitz, O.J., Hawlena, D. & Trussell, G.C. (2010) Predator control of ecosystem nutrient dynamics. Ecology Letters, 13, 1199–1209.

Schulte to Bühne, H. & Pettorelli, N. (2018) Better together: Integrating and fusing multispectral and radar satellite imagery to inform biodiversity monitoring, ecological research and conservation science. Methods in Ecology and Evolution, 9, 849–865.

Sebastián-González, E. et al. (2020) Network structure of vertebrate scavenger assemblages at the global scale: drivers and ecosystem functioning implications. Ecography, 43, 1143–1155.

Sella, G.F. et al. (2007) Observation of glacial isostatic adjustment in “stable” North America with GPS. Geophysical Research Letters, 34.

Smith, C.A.S., Smits, C.M.M. & Slough, B.G. (1992) Landform selection and soil modifications associated with Arctic fox (Alopex lagopus) den sites in Yukon Territory, Canada. Arctic and Alpine Research, 24, 324–328.

Smits, C.M.M., Smith, C.A.S. & Slough, B.G. (1988) Physical Characteristics of Arctic Fox (Alopex lagopus) Dens in Northern Yukon Territory, Canada. Arctic, 41, 12–16.

Speed, J.D.M. et al. (2019) Trophic interactions and abiotic factors drive functional and phylogenetic structure of vertebrate herbivore communities across the Arctic tundra biome. Ecography, 42, 1152–1163.

Tognelli, M.F. & Kelt, D.A. (2004) Analysis of determinants of mammalian species richness in South America using spatial autoregressive models. Ecography, 27, 427–436.

Turner, W. et al. (2015) Free and open-access satellite data are key to biodiversity conservation. Biological Conservation, 182, 173–176.

Valerio, F. et al. (2020) Predicting Microhabitat Suitability for an Endangered Small Mammal Using Sentinel-2 Data. Remote Sensing, 12.

van Moorter, B. et al. (2013) Understanding scales of movement: animals ride waves and ripples of environmental change. Journal of Animal Ecology, 82, 770–780.

Wood, S.N. (2011) Fast stable restricted maximum likelihood and marginal likelihood estimation of semiparametric generalized linear models. Journal of the Royal Statistical Society: Series B (Statistical Methodology), 73, 3–36.

Zupanc, A. (2017) Improving Cloud Detection with Machine Learning. https://medium.com/sentinel-hub/improving-cloud-detection-with-machine-learning-c09dc5dgfcx7cf13. Accessed 07 May 2022.

